# The universe is asymmetric, the mouse brain too

**DOI:** 10.1101/2023.09.01.555907

**Authors:** Alejandro Rivera-Olvera, Danielle J. Houwing, Jacob Ellegood, Shang Masifi, Stephany LL. Martina, Andrew Silberfeld, Olivier Pourquie, Jason P. Lerch, Clyde Francks, Judith R. Homberg, Sabrina van Heukelum, Joanes Grandjean

## Abstract

Hemispheric brain asymmetry is a basic organizational principle of the human brain and has been implicated in various psychiatric conditions, including autism spectrum disorder. Brain asymmetry is not a uniquely human feature and is observed in other species such as the mouse. Yet, asymmetry patterns are generally nuanced, and substantial sample sizes are required to detect these patterns. In this pre-registered study, we use a mouse dataset from the Province of Ontario Neurodevelopmental Network, which comprises structural MRI data from over 2000 mice, including genetic models for autism spectrum disorder, to reveal the scope and magnitude of hemispheric asymmetry in the mouse. Our findings demonstrate the presence of robust hemispheric asymmetry in the mouse brain, such as larger right hemispheric volumes towards the anterior pole and larger left hemispheric volumes toward the posterior pole, opposite to what has been shown in humans. This suggests the existence of species-specific traits. Further clustering analysis identified distinct asymmetry patterns in autism spectrum disorder models, a phenomenon that is also seen in atypically developing participants. Our study shows potential for the use of mouse models in studying the biological bases of typical and atypical brain asymmetry but also warrants caution as asymmetry patterns seem to differ between humans and mice.

## Main

Asymmetry is a basic human brain organizational principle, reflected in anatomical and functional differences between the left and right hemispheres ^1–3^. Lateralized brain development is already observed early on during fetal development ^4^, and perturbations in asymmetry have been associated with several psychiatric conditions, such as autism spectrum disorder ^5,6^. This suggests that specific asymmetry patterns are crucial for optimal brain functioning. Recently, various genes have been implicated through large-scale genetic association mapping ^7,8^, but the biological determinants of human brain asymmetry remain poorly understood.

Brain asymmetry is observed across the animal kingdom. There are however differences between species. For instance, the brain torque observed in humans (i.e. a larger right prefrontal lobe but larger left parietal-occipital lobe) has not been observed in non-human primates ^9–11^. In rodents, several studies show functional asymmetries, for example for the left vs. right hippocampus ^12^ and the left vs. right auditory cortex ^13^. Such functional asymmetries have also been observed in humans ^14,15^, suggesting the existence of similarities in asymmetry patterns across humans and rodents. Rodent literature on structural asymmetries is more sparse but hippocampal asymmetries, in terms of volumes ^16,17^, synaptic plasticity ^18^, and receptor distribution have been reported. Recent research suggests that structural asymmetry in rodents also further develops postnatally, especially affecting the hippocampus and the entorhinal cortex ^17^. As such, rodents appear to be suitable models for investigating the mechanisms underlying brain asymmetry. However, previous studies have been performed in rather small groups, ranging from 20 to a maximum of 100 animals ^16,17,19–21^, and as asymmetry patterns are generally nuanced, even within human populations, substantial sample sizes are required to describe this phenomenon accurately.

In this pre-registered study, we set out to examine hemispheric asymmetry in mice using the Province of Ontario Neurodevelopmental Network (POND) dataset which comprises structural MRI scans in both healthy controls (wild-type) and genetic models for autism spectrum disorders ^22^. We observe distributed asymmetry patterns in wild-type mice, albeit these corresponded to small to very small effects that require large samples to discern. Moreover, the asymmetry patterns only bear partial resemblance with those observed in humans, e.g. a rightward volumetric bias in the isocortex toward the anterior pole, whereas a leftward cortical thickness bias is reported in humans ^23^. In genetic models of autism spectrum disorder, we observe distinct widespread patterns of asymmetry relative to wild-type controls. Most notably, we observe clusters indicating more positive asymmetry indices across the isocortex in transgenic animals relative to wild-type in a cluster, and the opposite in another cluster.

We obtained 2300 Jacobian maps derived from the non-linear deformations needed to fit a template from the POND repository. First, we examined the asymmetry index from the relative Jacobians in n = 878 wild-type animals (n_female_ = 290, n_male_ = 533, n_n/a_ = 55). The asymmetry index was calculated using: *(Volume*_*Right*_ *-Volume*_*Left*_*) / (Volume*_*Right*_ *+ Volume*_*Left*_). The asymmetry index was derived from the region-of-interest volumes extracted from the DSURQE atlas (**Figure 1a**) ^24^. The DSURQE template is symmetrical, however, the label atlas is not, which can bias asymmetry index estimates. To account for this, we mirrored the right-hemisphere labels atlas onto the left hemisphere. Using the label atlas, we extracted the averaged Jacobians within each region of interest and multiplied it with the atlas region of interest volumes to obtain individual regional volumes. We then estimated Hedge’s g across the whole wild-type population (**Figure 1b**). We found both positive and negative Hedge’s g values (**Table S1**), e.g. in the striatum (g_wt > 0_ = 0.240 [0.146, 0.334]), cingulate cortex: area 24b’ (g_wt > 0_ = -0.194 [-0.288, -0.100]), indicating a rightward and leftward bias respectively. In all instances, the effect sizes were either very small (g < 0.2) or small (g ∼ 0.2), i.e. effects barely discerned by the naked eye ^25^. Because the effects are barely visible, we sought to address how reproducible they were. We resampled the wild-type groups into two groups over 500 iterations and estimated the correlation of the averaged asymmetry indices between the two groups (**Figure 1c**). We obtained a median correlation (r = 0.631) and therefore concluded that the asymmetry indices in wild-type animals are robust. Intriguingly, the rightward bias observed in the striatum (caudoputamen) is different from the observation in humans that reported a leftward bias in the putamen ^26,27^.

**Figure 1.**
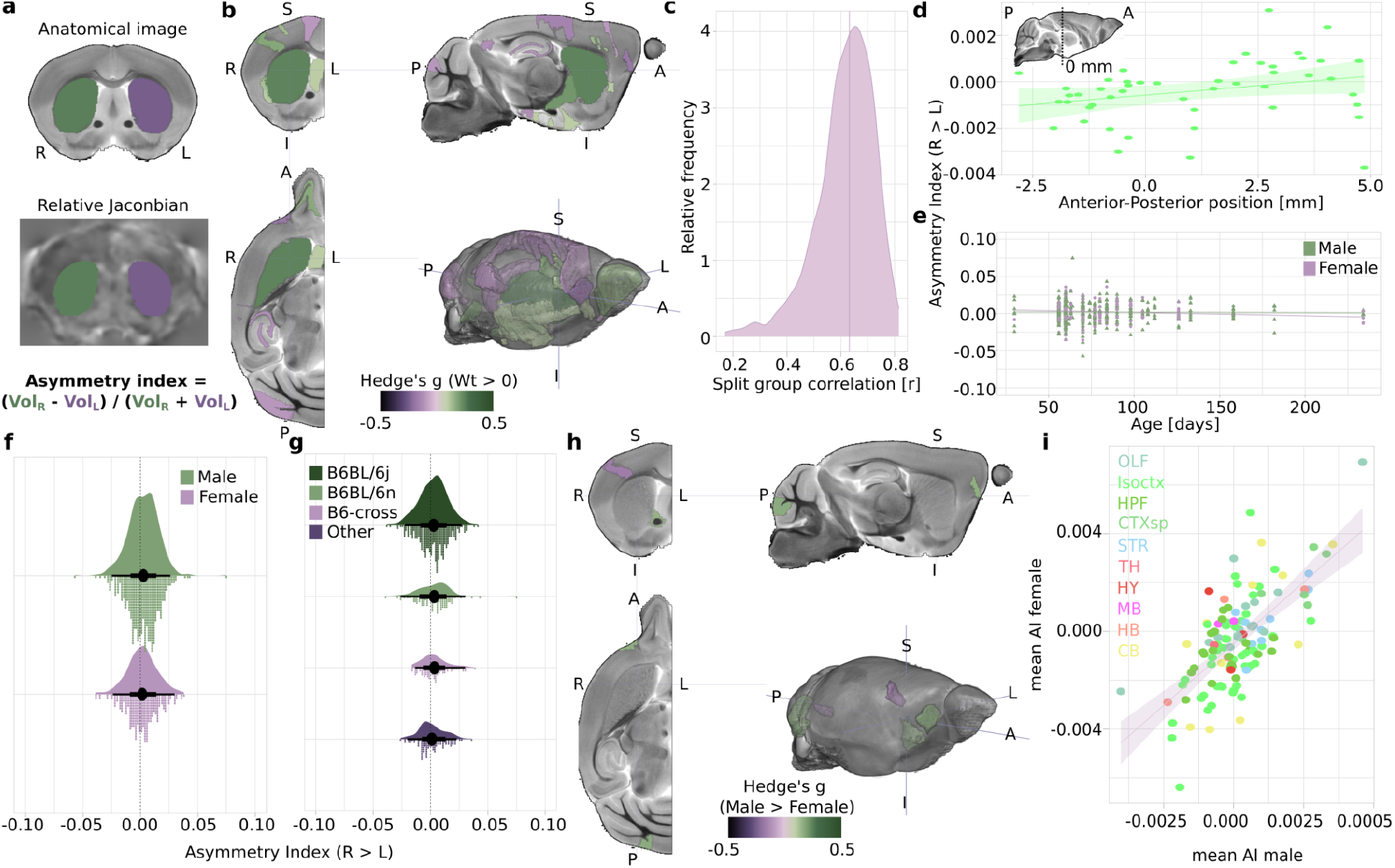
Asymmetry index in wild-type. **a)** Graphical representation of the asymmetry index, showing the anatomical template with region of interest labels for the striatum overlaid (top), an example relative Jacobian map with labels overlaid (bottom). **b)** Hedge’s g plot overlaid on the template for wild-type > 0 comparisons. The color code indicates thresholded Hedge’s g for both confidence intervals > or < 0 for positive or negative g values respectively. **c)** Distribution of Pearson’s correlation coefficients upon splitting the wild-type groups over 500 iterations. **d)** Average asymmetry index in wild-type animals as a function of the region-of-interest center of gravity along the Anterior-Posterior axis. The line and ribbon indicate the regression line and its 95% confidence intervals. The insert shows an outline of the brain with the origin position. **e)** Striatum asymmetry index in wild-type animals as a function of age. Dots indicate individual animals. The lines indicate the regression lines as a function of sex. **f)** Raincloud plots of the striatum asymmetry index in wild-type animals as a function of sex. Dots indicate individual animals. Black dots indicate the mean, thick and thin line intervals indicate the 66^th^ and 95^th^ percentiles. **g)** Raincloud plots of the striatum asymmetry index in wild-type animals as a function of sub-strains. **h)** Hedge’s g plot overlaid on the template for male > female comparisons in wild-type. **i)** Average asymmetry index in males and females with the Allen Brain Institute color code.

Still in the wild-type group, we sought to establish the presence of an asymmetry index pattern along the anterior-posterior axis. Indeed, this is one of the dominant features of hemispheric asymmetry in the human brain ^23^. Here, we restricted the analysis to the isocortical region of interest exclusively to compare with previous human studies. We found a small effect (**Figure 1d**, slope _AI ∼ Anterior-Posterior position_ = 1.64e^-4^ [-1.91e^-6^, 3.29e^-4^]). We concluded that there was a plausible effect along the anterior-posterior axis, with larger right hemispheric volumes towards the anterior pole and larger left hemispheric volumes toward the posterior pole. This is intriguing as a left > right asymmetry for cortical surface and thickness was instead observed in humans along the anterior-posterior axis ^7,23,28^.

Next, we describe the asymmetry index distribution in wild-type mice using the striatum as an example. For reference, the average striatum volume across wild-type animals was 6.848 ± 0.196 mm^3^ on the right hemisphere and 6.837 ± 0.196 mm^3^ on the left, or a 0.16% volume difference in the left hemisphere relative to the right. We could not conclude that there was an age effect in either females (**Figure 1e**, Slope_AI ∼ Age_ = -4.57e^-5^ [-1.01e^-4^, 9.25e^-6^]) or males (Slope_AI ∼ Age_ = -4.62e^-6^ [-4.46e^-4^, 3.54e^-5^]). There was a small bias towards positive (rightward) asymmetry index in both sexes (**Figure 1f**, n_females_ = 290, mean_females_ = 0.00171 ± 0.0132, n_males_ = 533, mean_males_ = 0.00267 ± 0.0131), as well as in all sub-strains included in the dataset (**Figure 1g**, n_C57BL/6J_ = 366, mean_C57BL/6J_ = 0.00264 ± 0.0126, n_C57BL/6N_ = 105, mean_C57BL6N_ = 0.00322 ± 0.0150, n_B6 crosses_ = 85, mean_B6 crosses_ = 0.00347 ± 0.0150, n_others_ = 109, mean_others_ = 0.00117 ± 0.0115). Taken together, we observe an asymmetry index bias evenly distributed across sexes and sub-strains. This brings added confidence beyond the confidence intervals of the effect size and the sub-sampling correlations that we are facing a plausible and consistent, albeit small, effect in wild-type animals.

Finally, we sought sex effects in wild-type animals across the whole brain. In humans, sex differences for asymmetry patterns are inconclusive and often show discrepancies in implicated brain regions or direction of effects ^1,26,29,30^. We found limited effects (**Table S2**), all very small, across the brain, notably in the bed nucleus of the stria terminalis (**Figure 1h**, n_female_ = 290, n_male_ = 533, g_male > female_ = 0.150 [0.00643, 0.293]). This is especially interesting as sex effects on functional activity of this region have been observed ^31,32^. When correlating the average asymmetry index between sexes, we found a large correlation (r = 0.671). We conclude from this analysis that hemispheric asymmetry sex differences are likely very small in mice and/or restricted to few areas. This is in line with observations in humans (n = 17,141), at least with regard to cortical thickness, where no regional sex effects were observed ^23^. There were instead small sex effects in striatal regions, e.g. rightwards shift in putamen asymmetry in males relative to females ^26^. In brief, our results do not fundamentally differ from the observations in humans with regard to sex.

The POND dataset consists of 63 studies based on transgenic mouse models for autism spectrum disorders, each containing wild-type and transgenic animals. We identified 87 plausible contrasts within these 63 studies, as some studies contain wild-type, heterozygotes, and homozygotes knock-outs, copy number variations such as microdeletion (df) or microduplication (dp), or other genetic alterations. Sample size varied between studies (range n = [3, 34]). We found a wide range of effects across the brain. For illustration purposes, we show the contrast outcomes from two lines, Chd8 (**Figure 2a-d**) and 16p11 (**Figure 2ef**), with the former coming from two datasets (“Nord” and “Basson”). To fully represent the contrast maps we present them un-thresholded. We observe large effects (g > 1) in several regions of interest. We find strikingly matching patterns in the wild-type > transgenic contrasts analysis of the Chd8 datasets and to a lesser extent in the 16p11 dataset, including positive g values across most of the isocortex, and negative g values in the ventral striatum and adjacent areas, and the retrohippocampal areas. This suggests that transgenic models of autism spectrum disorders may cluster into a few distinct observable asymmetry phenotypes.

**Figure 2.**
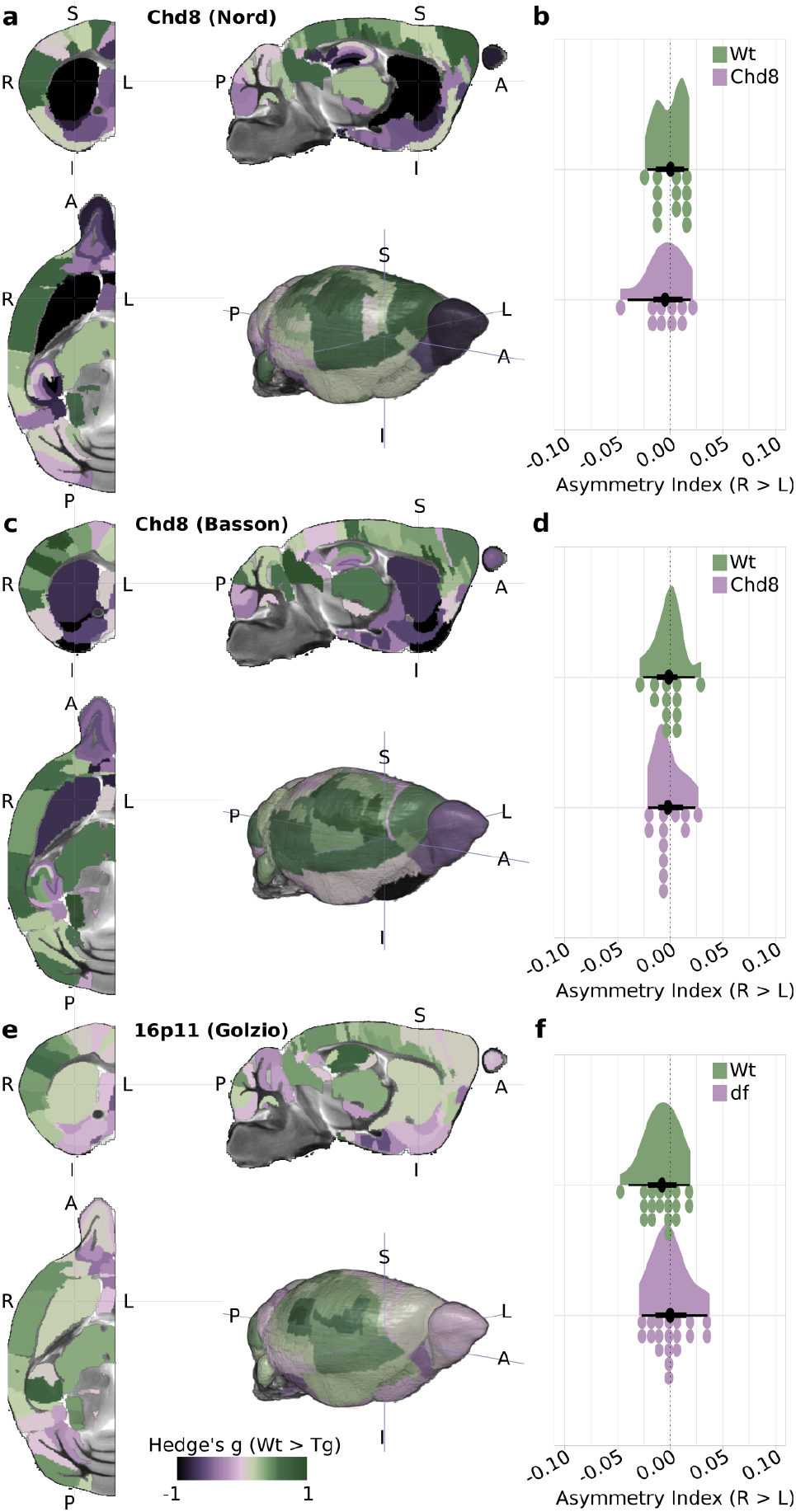
Selected model comparisons to wild-type controls. **a)** Hedge’s g plot overlaid on the template for wild-type > transgenic comparisons in Chd8 mice from the “Nord” dataset. The color code indicates unthresholded Hedge’s g. **b)** Raincloud plots of the striatum asymmetry index in wild-type (Wt, n = 12) and transgenic (Chd8, n = 10) Chd8 animals from the “Basson” dataset. Dots indicate individual animals. Black dots indicate the mean, thick and thin line intervals indicate the 66^th^ and 95^th^ percentiles. **c)** Hedge’s g plot overlaid on the template for wild-type > transgenic comparisons in Chd8 mice from the “Basson’’ dataset. **d)** Raincloud plots of the striatum asymmetry index in wild-type (Wt, n = 12) and transgenic (Chd8, n =12) Chd8 animals from the “Basson” dataset. **e)** Hedge’s g plot overlaid on the template for wild-type > transgenic (df) comparisons in 16p11 mice from the “Golzio” dataset. **f)** Raincloud plots of the striatum asymmetry index in wild-type (Wt, n = 18) and transgenic (df, n = 19) 16p11 animals from the “Golzio” dataset.

It is difficult to draw conclusions from individual contrasts as they are based on small samples ^33,34^. To overcome this, we hypothesized that, like brain volumes ^22^, the asymmetry index contrasts would cluster in these models. We ran a k-mean clustering algorithm on the asymmetry index contrasts x region-of-interest matrix across a range of k values (range = [1, 10]). We could not identify “elbows” in either the total within the sum of square or gap statistic plots (**Figure 3ab**), a common heuristic for selecting k values. Without these, we relied on a previous 3-cluster solution based on brain volumes as our heuristic for selecting k = 3 cluster here ^22^. Using this solution, we find plausible asymmetry index contrast outcome clusters (**Figure 3c, Table S3**). Firstly, the clusters show spatial organization. For instance, cluster #3 has generally positive g values in the isocortex regions of interest, while cluster #1 has negative values, and cluster #2 displays mixed g values. Secondly, the clusters gather contrasts from similar lines from different origins (e.g. Chd8 “Basson” and “Nord” in cluster #3). Thirdly, the clusters solution partially overlaps with that from Ellegood et al. 2015, for instance, cluster #1 contains NRXN1a (heterozygotes and homozygotes), SHANK3 (heterozygotes and homozygotes) similar to group #1 in Ellegood et al. To graphically represent the cluster solutions, we aggregated all the wild-types and transgenic for each cluster and estimated the cluster group differences. The spatial representations of the clusters recapitulate the observations in the matrice, namely cluster #1 consisting chiefly of negative g values in the isocortex, indicating more negative asymmetry index values in transgenic animals relative to wild-type (**Figure 3e**). Conversely, clusters #2 and #3 display chiefly positive g values in the lateral isocortex and in the striatum. Notably, the g value range in the clustered contrast analysis is lower than with individual contrasts, with all effects reported being either very small (g < 0.2) or small (g ∼ 0.2). For instance, in the striatum (**Figure 3d**), we find very small positive g values that pass our threshold criteria for cluster #2 (n_Wt_ = 738, n_Tg_ = 670, g_Wt > Tg_ = 0.115 [0.0421, 0.187]) and cluster #3 (n_Wt_ = 492, n_Tg_ = 365, g_Wt > Tg_ = 0.142 [0.0533, 0.231]), while cluster #1 did not (n_Wt_ = 234, n_Tg_ = 199, g_Wt > Tg_ = 0.104 [-0.0249, 0.233]). In sum, we observe biologically plausible clusters of asymmetry index contrasts in mouse models of autism spectrum disorder.

**Figure 3.**
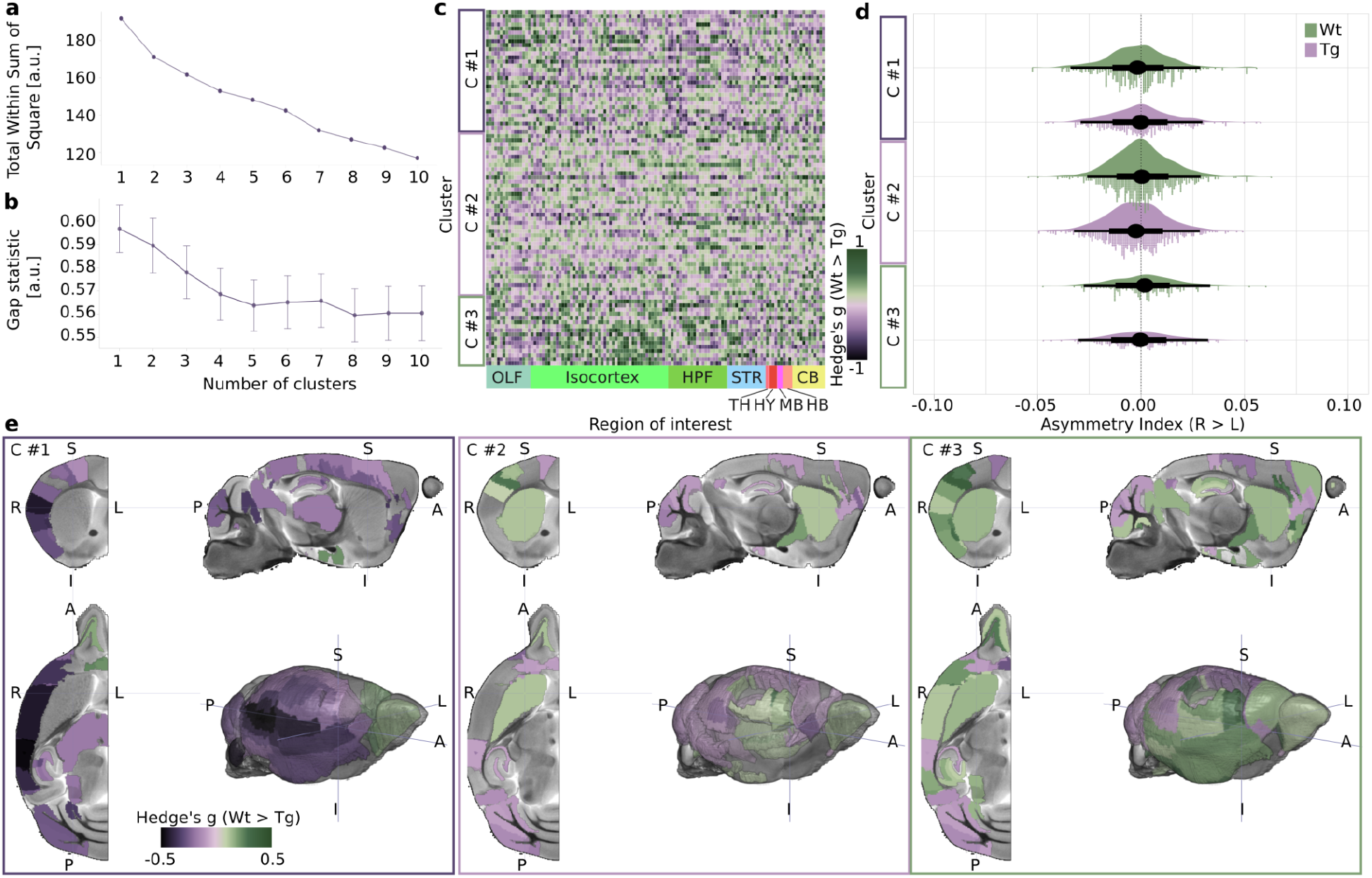
Clustering wild-type > transgenic contrasts. **a)** Total Within Sum of Square for different clustering solutions. **b)** Gap statistics for different clustering solutions. **c)** 3 cluster solution for wild-type > transgenic contrasts (y-axis) as a function of region-of-interest (x-axis). The region-of-interests are grouped and color-coded according to the Allen Brain Institute code. **d)** *Raincloud plots of the striatum asymmetry index in wild-type (Wt, n*_*cluster#1*_ *=* 234, *n*_*cluster#2*_ *=* 738, *n*_*cluster#3*_ *=* 492*) and transgenic (Tg, n*_*cluster#1*_ *=* 199, *n*_*cluster#2*_ *=* 670, *n*_*cluster#3*_ *=* 492*) animals as a function of clusters. Dots indicate individual animals. Black dots indicate the mean, thick and thin line intervals indicate the 66*^*th*^ *and 95*^*th*^ *percentiles*. ***e)*** *Hedge’s g plot overlaid on the template for wild-type > transgenic comparisons as a function of clusters. The color code indicates thresholded Hedge’s g for both confidence intervals > or < 0 for positive or negative g values respectively*.

These clusters consist of very small, but consistent effects. Notably, the observation in the striatum contrasts with what was reported in autism spectrum disorder participants relative to controls using the ENIGMA datasets ^35^, where an opposite very small effect size was observed in putamen volume (n_typical_ ∼ 800, n_atypical_ ∼ 500, Cohen’s d_typical > atypical_ = -0.12), denoting greater asymmetry index in atypical developing participants relative to typical developing participants ^27^. This difference is further reflected in typical developing participants where a leftward asymmetry bias was observed, relative to a rightward bias in mice here (**Figure 1b**). Further comparisons, in particular of the cortical region of interest, are harder to interpret as the analysis in Postema et al. relies on cortical surface and thickness rather than volumes as we do in this study. It should also be noted that the POND datasets are enriched in studies modeling syndromic autism, i.e. autism is accompanied by a host of other symptoms ^36,37^, while the ENIGMA contains both syndromic and idiopathic cases, i.e. autism is of unknown origin. This is expected to yield considerable differences in the outcomes between the mouse and human studies and should be considered a limitation when attempting to compare the two species. Nevertheless, the clusters identified here have the potential to explain phenotypic differences associated with syndromic autism and understand the biological context associated with the phenotype.

Our study demonstrates hemispheric asymmetry in mice and sets the stage to approach the biological context leading to this phenotype. The major limitations therein lie in the large sample size required to uncover the small to very small effects identified here. Indeed, small sample sizes generally lead to falsely inflated effects ^33,34^. We also observe this to some extent, as individual contrast maps display often large effects, while clustered contrast indicates very small to small effects exclusively. This likely implies that further biological investigations will rely either on large samples of wild-type animals or on genetic variations that may induce larger effects, such as microtubule-related genes (e.g. *TUBB3, TUBA1B, MAP2, MAPRE3*) identified to be associated with asymmetry outcomes in humans in the UK Biobank (n = 32,256) dataset ^8^. Microtubules form an essential component of the cytoskeleton, and the cytoskeleton has been shown to be involved in left-right axis determination of visceral organs in invertebrates and frogs ^38–40^. While the mechanisms of establishing brain asymmetry are currently unknown, mutations in microtubule-related genes such as TUBB3 induce severe phenotypes, characterized by cortical disorganization and axonal abnormalities ^41^. This suggests a role for microtubules in driving cortical organization, including the formation of brain asymmetry.

Another notable departure from human studies is the parameters used to estimate asymmetry indices. In cortical areas, studies in humans are commonly performed either on cortical thickness or surface ^23^, while we relied on volumes for both cortical and sub-cortical areas. The freesurfer toolbox allows for the seamless extraction of these parameters on human brains ^42^, however, the same toolbox has not been applied to the mouse. Due to a lack of equivalent toolkits in mice, cortical thickness analyses are rare ^43^. As cortical thickness and surface lead to vastly different outcomes, it is plausible that cortical volumes lead to further different outcomes. Hence, discrepancies between this study and studies in humans can be related to different parameters. Still, this does not suffice to explain all the discrepancies. For instance, in the striatum, we report a rightward bias in the caudoputamen volumes, while there is a leftward bias for the putamen in humans. Beyond looking at structural markers, there is also evidence for functional asymmetries, including in autism spectrum disorders, for instance using functional gradients ^44^, that would be interesting to investigate in corresponding animal models ^45,46^. This could unlock additional comparisons between species and help to bring an understanding of functional asymmetries.

In conclusion, we observe robust, albeit small to very small, hemispheric brain asymmetry in wild-type mice. These patterns present an anterior-posterior axis organization, with rightward biases toward the anterior pole. When examining transgenic models of autism spectrum disorders, we find biologically plausible clusters that relate to previous clustering solutions in these models ^22^. While the observations we obtained appear robust along several criteria, the outcomes depart from major observations in humans ^23,26,27^. This suggests that aspects of brain hemisphere asymmetry are distinctly found in humans. This is not fully inconsistent with the known biology of brain asymmetry, as aspects of asymmetry could not be highlighted in non-human primates either ^9–11^. Still, these observations in mice are crucial in understanding the phenomena due to the many available transgenic lines that allow for the testing in experimental conditions of relevant genetic variations, as well as the detailed description of the contribution of early developmental stages to the formation of these patterns.

## Method

### Deviations from pre-registration

This study was pre-registered (https://osf.io/bufr9) ^47^. Our project deviated from the pre-registration as we did not perform group variance testing. This is justified because the sample size for individual study groups was too small and the variance test is prone to false outcomes in small samples. We also updated the preregistration prior to the analysis onset to simplify the statistical models to better reflect the structure of the data. This is justified because sex and other co-variates are not distributed orthogonally from the studies and therefore cannot be easily modeled using multiple regression models that encompass all studies together.

### Data and code availability

The raw data is available from the POND repository (https://portal.conp.ca/dataset?id=projects/braincode_Mouse_Image) following access requests addressed to the data owners (JE, JPL). The processed data and all the steps to replicate the analysis are available under the terms of the Apache-2 license (www.github.com/grandjeanlab/mouse_asymmetry). To run the code in a similar environment as the one used for the analysis, the recipe for the generation of a reproducible Apptainer/Singularity container solution for GNU/Linux systems including all the analysis dependencies is available on the same GitHub repository ^48^.

### Preprocessing

Raw images were preprocessed in a previous work ^22^. Absolute and relative Jacobian images obtained from the linear and non-linear registrations to the DSURQE mouse brain template were imported into *Python3* (3.10.11) using the *Nibabel* package (5.1.0) ^38^. The averaged Jacobians were extracted using the DSURQE label atlas using the *masker* function implemented in *Nilearn* (0.10.1) ^39^. The DSURQE label atlas right hemisphere region of interests were mirrored onto the left hemisphere to ensure that paired region of interests have the same nominal volumes in the template. The averaged Jacobians were multiplied with the region of interest volumes to obtain individual region of interest volumes. The asymmetry index was estimated with the formula below.

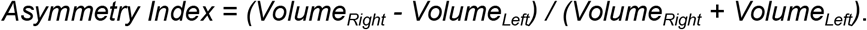

### Analysis and statistics

Asymmetry indices were imported into *R* (4.1.2) with the *tidyverse* package suit (1.3.1) ^51^. Descriptive statistics are provided as group mean ± 1 standard deviation. In an effort to move beyond p-value-based inferences ^52^, we rely on effect size and their 95% confidence intervals for inferences. Specifically, we thresholded effect size if both 2.5% and 97.5% confidence intervals were either > 0 for positive effect size or < 0 for negative effect size. We opted for Hedge’s g as the effect size. This is justified because i) we performed one or two group comparisons in the absence of co-variates exclusively, and ii) because comparisons in transgenic animal models rely on small samples for which the Hedge’s g effect size is a better indicator. We use the Cohen 1988 rules for effect size interpretations, namely: very small, g < 0.2; small, g > 0.2; medium, g > 0.5, large, g > 0.8 ^25^. Effect size and their 95% confidence intervals were estimated using the *effectsize* package (0.6.0.1) ^53^ and reported as g values [2.5^th^ interval, 97.5^th^ interval] for two sample comparisons. K-mean clusters are estimated using the *cluster* (2.1.4) and *factoextra* (1.0.7) packages ^54,55^. Plots are generated using *ggplot2* (3.4.2) together with *ggdist* (3.3.0) for Raincloud plots ^56,57^. Brain maps are rendered using *MRIcroGL* (v1.2.20220720). The color theme is “Cassatt2” from the *MetBrewer* package (0.2.0) ^58^.

## Acknowledgment

This project was supported by the Horizon Europe programs CANDY under grant agreement nos. 847818 to JG and JRH.

## CRediT author contribution

**Alejandro Rivera-Olvera** - Conceptualization, Formal Analysis, Writing – Original Draft**; Danielle J. Houwing** - Conceptualization, Writing – Review & Editing**; Jacob Ellegood** - Data Curation, Writing – Review & Editing**; Shang Masifi** - Writing – Review & Editing**; Stephany LL. Martina** - Writing – Review & Editing**; Andrew Silberfeld** - Conceptualization, Writing – Review & Editing**; Olivier Pourquie -** Conceptualization, Writing – Review & Editing**; Jason P. Lerch** - Funding Acquisition, Writing – Review & Editing**; Clyde Francks** - Conceptualization, Supervision, Writing – Review & Editing**; Judith R. Homberg** - Funding Acquisition, Supervision, Writing – Review & Editing**; Sabrina van Heukelum** - Conceptualization, Formal Analysis, Supervision, Writing – Original Draft**; Joanes Grandjean** - Conceptualization, Formal Analysis, Funding Acquisition, Software, Supervision, Visualization, Writing – Original Draft

## Supplementary material

**Table S1.** Contrast outcome for AI_wild-type > 0_.

| Region of interest | Hedge's g [2.5<sup>th</sup> CI, 97.5<sup>th</sup> CI] |
| --- | --- |
| striatum | 0.240 [0.146, 0.334] |
| fastigial nucleus | 0.176 [0.083, 0.270] |
| Cingulate cortex: area 24a' | -0.108 [-0.201, -0.014] |
| Cingulate cortex: area 24b' | -0.194 [-0.288, -.010] |
| Cingulate cortex: area 29b | -0.140 [-0.234, -0.047] |
| Cingulate cortex: area 29c | -0.202 [-0.296, -0.109] |
| Cingulate cortex: area 30 | -0.12 [-0.213, -0.260] |
| globus pallidus | 0.205 [0.111, 0.298] |
| Primary auditory cortex | -0.113 [-0.207, -0.020] |
| Secondary auditory cortex: ventral area | -0.122 [-0.216, -0.029] |
| amygdala | 0.095 [0.002, 0.189] |
| Clastrum: dorsal part | 0.137 [0.044, 0.231] |
| Dorsal nucleus of the endopiriform | 0.101 [0.008, 0.195] |
| Dorsolateral orbital cortex | -0.188 [-0.281, -0.094] |
| Frontal cortex: area 3 | 0.137 [0.044, 0.231] |
| Secondary motor cortex | -0.098 [-0.192, -0.005] |
| flocculus (FL) | 0.200 [0.106, 0.293] |
| superior olivary complex | 0.187 [0.094, 0.281] |
| Primary somatosensory cortex: hindlimb region | -0.154 [-0.247, -0.06] |
| Primary somatosensory cortex: jaw region | 0.175 [0.081, 0.269] |
| Clastrum: ventral part | 0.122 [0.028, 0.215] |
| Ventral nucleus of the endopiriform claustrum | 0.131 [0.037, 0.224] |
| Ventral tenia tecta | -0.105 [-0.199, -0.012] |
| cuneate nucleus | -0.169 [-0.263, -0.075] |
| crus 2: ansiform lobule (lobule 7) | -0.103 [-0.197, -0.01] |
| lateral septum | 0.097 [0.003, 0.190] |
| LMol | -0.116 [-0.209, -0.022] |
| CA3Py Inner | -0.115 [-0.208, -0.021] |
| CA30r | -0.125 [-0.218, -0.031] |
| PoDG | -0.110 [-0.203, -0.016] |
| Olfactory bulb: internal plexiform layer | 0.126 [0.033, 0.22] |
| Olfactory bulb: granule cell layer | 0.172 [0.078, 0.265] |
| Accessory olfactory bulb: granule cell layer | 0.272 [0.178, 0.366] |

**Table S2.** Contrast outcome for AI_male > female_.

| Region of interest | Hedge's g [2.5<sup>th</sup> CI, 97.5<sup>th</sup> CI] |
| --- | --- |
| Fastigial nucleus | -0.171 [-0.314, -0.0278] |
| Dorsolateral orbital cortex | 0.159 [0.0158, 0.302] |
| Paramedian lobule (lobule 7) | 0.166 [0.0225, 0.309] |
| Primary somatosensory cortex: jaw region | -0.172 [-0.315, -0.0291] |
| Bed nucleus of stria terminalis | 0.150 [0.00643, 0.293] |

**Table S3.** Cluster allocation per contrast.

| cluster | study | Wt group | Wt n | Tg group | Tg n |
| --- | --- | --- | --- | --- | --- |
| 1 | 15q_wMat | WT | 8 | MDp | 9 |
| 1 | 22q11 | WT | 10 | DEL | 10 |
| 1 | BTBR | WT | 12 | BTBR | 12 |
| 1 | BTBR | FVB | 12 | BTBR | 12 |
| 1 | CNTNAP2 | WT | 9 | KO | 9 |
| 1 | FMR1_FVB | WT | 10 | KO | 10 |
| 1 | Gtf2i | WT | 20 | YT | 20 |
| 1 | Gtf2i | WT | 20 | XS | 18 |
| 1 | MAR | WT | 23 | MAR | 22 |
| 1 | NF1 | WT | 9 | HET | 10 |
| 1 | NL3 | WT | 8 | KO | 8 |
| 1 | NRXN1a | WT | 10 | HET | 13 |
| 1 | NRXN1a | WT | 10 | KO | 9 |
| 1 | SERT_KI | WT | 20 | KI | 20 |
| 1 | SHANK3 | WT | 10 | HET | 11 |
| 1 | SHANK3 | WT | 10 | KO | 9 |
| 1 | Ube3a | WT | 17 | HET | 15 |
| 1 | CHD7_En1Cre | En1Cre | 12 | En1CreChd7f+ | 10 |
| 1 | 16p11_Golzio | WT | 12 | dp | 9 |
| 1 | Kctd13_Golzio | WT | 19 | HET | 21 |
| 1 | Nphp1 | WT | 7 | Tg | 9 |
| 1 | Lat_WT | WT | 9 | HET | 12 |
| 1 | Lat_WT | WT | 9 | HOM | 9 |
| 1 | Tbr1_WT | WT | 9 | HET | 9 |
| 1 | CMMR_New | WT | 38 | rab39b | 14 |
| 1 | CMMR_New | WT | 38 | ranbp17 | 11 |
| 1 | CMMR_New | WT | 38 | upf3b | 13 |
| 1 | CMMR_New | WT | 38 | ypel2 | 9 |
| 1 | CMMR_New | WT | 38 | l2hgdh | 15 |
| 1 | GSK3 | WT | 7 | Beta | 7 |
| 2 | 15q | WT | 10 | pDp | 10 |
| 2 | 16p11 | WT | 9 | dp | 11 |
| 2 | AndR | WT | 22 | 12Q | 17 |
| 2 | AndR | WT | 22 | 48Q_Het | 10 |
| 2 | Arhgef6 | WT | 32 | Arhgef6 | 16 |
| 2 | BALBC | WT | 9 | BALBC | 12 |
| 2 | Dpyd | WT | 11 | HET | 22 |
| 2 | Dpyd | WT | 11 | HOM | 12 |
| 2 | Dvl1 | WT | 21 | KO | 26 |
| 2 | En2 | WT | 9 | KO | 9 |
| 2 | ITGB3 | WT | 11 | KO | 11 |
| 2 | Magel2 | WT | 6 | MUT | 6 |
| 2 | NL1 | WT | 26 | KO | 27 |
| 2 | SERT_KI_129 | WT | 21 | KI | 21 |
| 2 | TSC1 | WT | 26 | MUT | 33 |
| 2 | Turner | WT | 23 | XO | 10 |
| 2 | CHD7_Het | WT | 13 | Chd7gt+ | 13 |
| 2 | CHD7_En1Cre | En1Cre | 12 | En1CreChd7ff | 10 |
| 2 | Cacnb2 | WT | 11 | Cacnb2 | 15 |
| 2 | Scn1a | WT | 3 | HET | 10 |
| 2 | Kctd13_Golzio | WT | 19 | HOM | 19 |
| 2 | Chd8_Kim | WT | 30 | Chd8 | 30 |
| 2 | Kctd13 | WT | 23 | HET | 20 |
| 2 | Kctd13 | WT | 23 | KO | 23 |
| 2 | Ube3a_Dup | WT | 13 | tTA | 11 |
| 2 | Ube3a_Dup | WT | 13 | Dup15 | 12 |
| 2 | Taok2 | WT | 16 | HET | 13 |
| 2 | Taok2 | WT | 16 | KO | 23 |
| 2 | Cul3 | WT | 18 | HET | 20 |
| 2 | tcdd | WT | 21 | TCDD | 34 |
| 2 | latXkctd13 | WT | 19 | Kctd13Het_LatHet | 22 |
| 2 | VPA | WT | 24 | VPA | 30 |
| 2 | Syngap1 | WT | 16 | HET | 18 |
| 2 | CMMR_New | WT | 38 | otc | 10 |
| 2 | CMMR_New | WT | 38 | pah | 15 |
| 2 | CMMR_New | WT | 38 | katnal2 | 14 |
| 2 | CMMR_New | WT | 38 | nexmif | 19 |
| 2 | GSK3 | WT | 7 | Alpha | 6 |
| 2 | Itsn | WT | 10 | itsn1 | 15 |
| 2 | Itsn | WT | 10 | itsn2 | 15 |
| 3 | 16p11 | WT | 9 | df | 11 |
| 3 | AndR | WT | 22 | 12Q_Het | 11 |
| 3 | AndR | WT | 22 | 21Q_Het | 7 |
| 3 | AndR | WT | 22 | 21Q | 12 |
| 3 | AndR | WT | 22 | 48Q | 15 |
| 3 | Arid1b | WT | 1 | Arid1b | 13 |
| 3 | Chd8_Basson | WT | 12 | Chd8 | 12 |
| 3 | Chd8_Nord | WT | 18 | Chd8 | 19 |
| 3 | FMR1 | WT | 8 | KO | 8 |
| 3 | SERT_KO | WT | 10 | KO | 20 |
| 3 | 16p11_Golzio | WT | 12 | df | 10 |
| 3 | 16p11_Golzio | WT | 12 | dfdp | 7 |
| 3 | Ube3a_Dup | WT | 13 | TRE | 10 |
| 3 | Setd5 | WT | 12 | HET | 11 |
| 3 | Itsn | WT | 10 | DKO | 3 |
| 3 | Mthfr | WT | 29 | HET | 30 |

## References

1. Toga, A. W. & Thompson, P. M. Mapping brain asymmetry. Nat. Rev. Neurosci. 4, 37–48 (2003).

2. Lin, S.-Y. & Burdine, R. D. Brain Asymmetry: Switching from Left to Right. Curr. Biol. 15, R343–R345 (2005).

3. Zhou, D., Lebel, C., Evans, A. & Beaulieu, C. Cortical thickness asymmetry from childhood to older adulthood. NeuroImage 83, 66–74 (2013).

4. Kasprian, G. et al. The prenatal origin of hemispheric asymmetry: an in utero neuroimaging study. Cereb. Cortex N. Y. N 1991 21, 1076–1083 (2011).

5. Oertel, V. et al. Reduced laterality as a trait marker of schizophrenia--evidence from structural and functional neuroimaging. J. Neurosci. Off. J. Soc. Neurosci. 30, 2289–2299 (2010).

6. Herbert, M. R. et al. Abnormal asymmetry in language association cortex in autism. Ann. Neurol. 52, 588–596 (2002).

7. Sha, Z. et al. Handedness and its genetic influences are associated with structural asymmetries of the cerebral cortex in 31,864 individuals. Proc. Natl. Acad. Sci. 118, e2113095118 (2021).

8. Sha, Z. et al. The genetic architecture of structural left–right asymmetry of the human brain. Nat. Hum. Behav. 5, 1226–1239 (2021).

9. Xiang, L., Crow, T. J., Hopkins, W. D., Gong, Q. & Roberts, N. Human torque is not present in chimpanzee brain. NeuroImage 165, 285–293 (2018).

10. Xiang, L., Crow, T. & Roberts, N. Cerebral torque is human specific and unrelated to brain size. Brain Struct. Funct. 224, 1141–1150 (2019).

11. Neubauer, S., Gunz, P., Scott, N. A., Hublin, J.-J. & Mitteroecker, P. Evolution of brain lateralization: A shared hominid pattern of endocranial asymmetry is much more variable in humans than in great apes. Sci. Adv. 6, eaax9935 (2020).

12. Jordan, J. T. The rodent hippocampus as a bilateral structure: A review of hemispheric lateralization. Hippocampus 30, 278–292 (2020).

13. Calhoun, G., Chen, C.-T. & Kanold, P. O. Bilateral widefield calcium imaging reveals circuit asymmetries and lateralized functional activation of the mouse auditory cortex. Proc. Natl. Acad. Sci. 120, e2219340120 (2023).

14. Spiers, H. J. et al. Unilateral temporal lobectomy patients show lateralized topographical and episodic memory deficits in a virtual town. Brain J. Neurol. 124, 2476–2489 (2001).

15. Devlin, J. T. et al. Functional Asymmetry for Auditory Processing in Human Primary Auditory Cortex. J. Neurosci. 23, 11516–11522 (2003).

16. Spring, S., Lerch, J. P., Wetzel, M. K., Evans, A. C. & Henkelman, R. M. Cerebral asymmetries in 12-week-old C57Bl/6J mice measured by magnetic resonance imaging. NeuroImage 50, 409–415 (2010).

17. Zeng, C. et al. Asymmetry of brain development in adolescent rats studied by 3.0 T magnetic resonance imaging. Neuroreport (2023) doi:10.1097/WNR.0000000000001943.

18. Shipton, O. A. et al. Left-right dissociation of hippocampal memory processes in mice. Proc. Natl. Acad. Sci. U. S. A. 111, 15238–15243 (2014).

19. Diamond, M. C., Dowling, G. A. & Johnson, R. E. Morphologic cerebral cortical asymmetry in male and female rats. Exp. Neurol. 71, 261–268 (1981).

20. Diamond, M. C., Johnson, R. E., Young, D. & Singh, S. S. Age-related morphologic differences in the rat cerebral cortex and hippocampus: male-female; right-left. Exp. Neurol. 81, 1–13 (1983).

21. Kolb, B., Sutherland, R. J., Nonneman, A. J. & Whishaw, I. Q. Asymmetry in the cerebral hemispheres of the rat, mouse, rabbit, and cat: the right hemisphere is larger. Exp. Neurol. 78, 348–359 (1982).

22. Ellegood, J. et al. Clustering autism - using neuroanatomical differences in 26 mouse models to gain insight into the heterogeneity. Mol. Psychiatry 20, 118–125 (2015).

23. Kong, X.-Z. et al. Mapping cortical brain asymmetry in 17,141 healthy individuals worldwide via the ENIGMA Consortium. Proc. Natl. Acad. Sci. U. S. A. 115, E5154–E5163 (2018).

24. Dorr, A. E., Lerch, J. P., Spring, S., Kabani, N. & Henkelman, R. M. High resolution three-dimensional brain atlas using an average magnetic resonance image of 40 adult C57Bl/6J mice. NeuroImage 42, 60–69 (2008).

25. Cohen, J. Statistical Power Analysis for the Behavioral Sciences. (Routledge, 1988). doi:10.4324/9780203771587.

26. Guadalupe, T. et al. Human subcortical brain asymmetries in 15,847 people worldwide reveal effects of age and sex. Brain Imaging Behav. 11, 1497–1514 (2017).

27. Postema, M. C. et al. Altered structural brain asymmetry in autism spectrum disorder in a study of 54 datasets. Nat. Commun. 10, 4958 (2019).

28. Roe, J. M. et al. Tracing the development and lifespan change of population-level structural asymmetry in the cerebral cortex. eLife 12, e84685.

29. Kurth, F., Thompson, P. M. & Luders, E. Investigating the Differential Contributions of Sex and Brain Size to Gray Matter Asymmetry. Cortex J. Devoted Study Nerv. Syst. Behav. 99, 235–242 (2018).

30. Luders, E., Gaser, C., Jancke, L. & Schlaug, G. A voxel-based approach to gray matter asymmetries. NeuroImage 22, 656–664 (2004).

31. Smithers, H. E., Terry, J. R., Brown, J. T. & Randall, A. D. Sex-associated differences in excitability within the bed nucleus of the stria terminalis are reflective of cell-type. Neurobiol. Stress 10, 100143 (2019).

32. Vantrease, J. E. et al. Sex differences in the Activity of Basolateral Amygdalar Neurons that Project to the Bed Nucleus of the Stria Terminalis and their Role in Anticipatory Anxiety. J. Neurosci. Off. J. Soc. Neurosci. 42, 4488–4504 (2022).

33. Button, K. S. et al. Power failure: why small sample size undermines the reliability of neuroscience. Nat. Rev. Neurosci. 14, 365–376 (2013).

34. Ioannidis, J. P. A. Why Most Discovered True Associations Are Inflated. Epidemiology 19, 640 (2008).

35. van Rooij, D. et al. Cortical and Subcortical Brain Morphometry Differences Between Patients With Autism Spectrum Disorder and Healthy Individuals Across the Lifespan: Results From the ENIGMA ASD Working Group. Am. J. Psychiatry 175, 359–369 (2018).

36. Ziats, C. A., Patterson, W. G. & Friez, M. Syndromic Autism Revisited: Review of the Literature and Lessons Learned. Pediatr. Neurol. 114, 21–25 (2021).

37. Molloy, C. J. et al. Bridging the translational gap: what can synaptopathies tell us about autism? Front. Mol. Neurosci. 16, 1191323 (2023).

38. Tee, Y. H. et al. Cellular chirality arising from the self-organization of the actin cytoskeleton. Nat. Cell Biol. 17, 445–457 (2015).

39. Davison, A. et al. Formin Is Associated with Left-Right Asymmetry in the Pond Snail and the Frog. Curr. Biol. CB 26, 654–660 (2016).

40. Lobikin, M. et al. Early, nonciliary role for microtubule proteins in left-right patterning is conserved across kingdoms. Proc. Natl. Acad. Sci. U. S. A. 109, 12586–12591 (2012).

41. Poirier, K. et al. Mutations in the neuronal ß-tubulin subunit TUBB3 result in malformation of cortical development and neuronal migration defects. Hum. Mol. Genet. 19, 4462–4473 (2010).

42. Dale, A. M., Fischl, B. & Sereno, M. I. Cortical surface-based analysis. I. Segmentation and surface reconstruction. NeuroImage 9, 179–194 (1999).

43. Lerch, J. P. et al. Cortical thickness measured from MRI in the YAC128 mouse model of Huntington’s disease. NeuroImage 41, 243–251 (2008).

44. Wan, B. et al. Diverging asymmetry of intrinsic functional organization in autism. Mol. Psychiatry 1–11 (2023) doi:10.1038/s41380-023-02220-x.

45. Huntenburg, J. M., Yeow, L. Y., Mandino, F. & Grandjean, J. Gradients of functional connectivity in the mouse cortex reflect neocortical evolution. NeuroImage 225, 117528 (2021).

46. Zerbi, V. et al. Brain mapping across 16 autism mouse models reveals a spectrum of functional connectivity subtypes. Mol. Psychiatry 26, 7610–7620 (2021).

47. Grandjean, J. Brain asymmetry in mouse models of autism spectrum disorder. (2022) doi:10.17605/OSF.IO/BUFR9.

48. Kurtzer, G. M. et al. hpcng/singularity: Singularity 3.7.3. (2021) doi:10.5281/zenodo.4667718.49.

49. Brett, M. et al. nipy/nibabel: 5.1.0. (2023) doi:10.5281/zenodo.7795644.

50. Chamma, A. et al. nilearn. (2023).

51. Wickham, H. et al. Welcome to the tidyverse. J. Open Source Softw. 4, 1686 (2019).

52. Ho, J., Tumkaya, T., Aryal, S., Choi, H. & Claridge-Chang, A. Moving beyond P values: data analysis with estimation graphics. Nat. Methods 16, 565–566 (2019).

53. Ben-Shachar, M. S., Lüdecke, D. & Makowski, D. effectsize: Estimation of Effect Size Indices and Standardized Parameters. J. Open Source Softw. 5, 2815 (2020).

54. Kassambara, A. & Mundt, F. factoextra: Extract and Visualize the Results of Multivariate Data Analyses. (2020).

55. Maechler, M., Rousseeuw, P., Struyf, A., Hubert, M. & Hornik, K. cluster: Cluster Analysis Basics and Extensions. (2022).

56. Wickham, H. ggplot2: Elegant Graphics for Data Analysis. (Springer-Verlag New York, 2016).

57. Kay, M. ggdist: Visualizations of Distributions and Uncertainty. doi:10.5281/zenodo.3879620.

58. Mills, B. R. MetBrewer: Color Palettes Inspired by Works at the Metropolitan Museum of Art. (2022).

